# The non-homologous end joining factor Ku orchestrates replication fork resection and fine-tunes Rad51-mediated fork restart

**DOI:** 10.1101/162677

**Authors:** Ana Teixeira-Silva, Anissia Ait Saada, Ismail Iraqui, Marina Charlotte Nocente, Karine Fréon, Julien Hardy, Sarah Lambert

## Abstract

Replication requires Homologous Recombination (HR) to stabilize and restart terminally-arrested forks. HR-mediated fork processing requires single stranded DNA (ssDNA) gaps and not necessarily Double Strand Breaks. We used genetic and molecular assays to investigate fork-resection and restart at dysfunctional, unbroken forks in *Schizosaccharomyces pombe*. We found that fork-resection is a two-step process coordinated by the non-homologous end joining factor Ku. An initial resection mediated by MRN/Ctp1 removes Ku from terminally-arrested forks, generating ~ 110 bp sized gaps obligatory for subsequent Exo1-mediated long-range resection and replication restart. The lack of Ku results in slower fork restart, excessive resection, and impaired RPA recruitment. We propose that terminally-arrested forks undergo fork reversal, providing a single DNA end for Ku binding which primes RPA-coated ssDNA. We uncover an unprecedented role for Ku in orchestrating resection of unbroken forks and in fine-tuning HR-mediated replication restart.

- Ku orchestrates a two-steps DNA end-resection of terminally-arrested and unbroken forks
- MRN/Ctp1 removes Ku from terminally-arrested forks to initiate fork-resection
- a ~110 bp sized ssDNA gap is sufficient and necessary to promote fork restart.
- The lack of Ku decreases ssDNA RPA-coating, and slows down replication fork restart.

## Introduction

At each cell division, ensuring the correct duplication and segregation of the genetic material is crucial for maintaining genome integrity. Although mutational events contribute to genome evolution, DNA lesions trigger genome instability often associated with human diseases such as cancer, genomic disorders, aging and neurological dysfunctions^1^.

DNA replication constitutes a major peril for genome stability. This is particularly evident in the case of oncogene-induced proliferation, which results in replication stress and faulty genome duplication, contributing to acquire genetic instability in neoplasic lesions^2,3^. A key feature of replication stress is the alteration of replication fork progression by numerous replication fork barriers (RFBs) that threaten faithful DNA duplication^4^. These hurdles are both intrinsic and extrinsic to the cell. Among them are non-histone protein-DNA complexes, the clashing of replication with transcription machineries, DNA secondary structures, damaged DNA, and nucleotide depletion.

RFBs impact replisomes’ functionality, and may result in replication forks stalling, requiring stabilization by the S-phase checkpoint to ensure DNA synthesis resumption^5^. RFBs may also result in dysfunctional and terminally-arrested forks, which lack their replication-competent state, and necessitate additional mechanisms to resume DNA synthesis. Through nucleolytic cleavage, terminally-arrested forks are converted into broken forks, exhibiting one ended DSB^6^. A DNA nick directly converts an active fork into a broken fork, accompanied with a loss of some replisome components^7^. Forks lacking their replication competence, and thus terminally-arrested, are often referred to as collapsed forks, whether broken or not.

To counteract deleterious outcomes of replication stress, cells have evolved the Homologous Recombination (HR) pathway which ensures the repair of DSBs and secures DNA replication^8^. HR is initiated by the loading of the recombinase Rad51 onto ssDNA, with the assistance of mediators such as yeast Rad52 and mammalian BRCA2. The Rad51 filament then promotes homology search and strand invasion with an intact homologous DNA template. HR allows the repair of forks exhibiting one ended DSBs likely through Break Induced Replication^9-12^. However, growing evidences point out that DSBs are not a pre-requirement neither for the recruitment of HR factors at dysfunctional forks nor for their restart^13-17^.

DSBs are also repaired by the non-homologous end joining (NHEJ) pathway which promotes the direct ligation of DNA ends with limited or no end-resection^18^. While HR is active in S and G2-phase, NHEJ is active throughout the cell cycle. A key component of NHEJ is the heterodimer Ku composed of two subunits, Ku70 and Ku80, both required for the stability of the complex. Yeast Ku binds dsDNA ends, inhibits end-resection, and allows the ligation of DSBs trough the recruitment of Ligase 4^19-21^. Interestingly, Ku is also involved in the repair of replication-born DSBs, where it limits end-resection^21-24^ and yeast Ku acts as a backup to promote cell survival upon replication stress^25,26^.

In most eukaryotes, DSBs resection is a two-step process^27,28^. An initial 5’ to 3’ nucleolytic processing, limited to the vicinity of the DNA end, is mediated by the MRN (Mre11/Rad50/Nbs1) complex which binds DSBs as an early sensor^29^. MRN then recruits Ctp1, a protein reported to share a conserved C-terminal domain with the mammalian nuclease CtIP and with *Saccharomyces cerevisiae* Sae2^30,31^. The endo- and exo-nuclease activities of Mre11, stimulated by Sae2/Ctp1, are not strictly required to DNA end resection at “clean” DSBs, but are critical to process “dirty” ends (generated by gamma-irradiation) or blocked ends such as meiotic DSBs at which Spo11 remains covalently attached to the DNA^19,32-35^. In yeasts, the nuclease activity of Mre11 and the Ctp1-dependent clipping function are required for Ku removal from DSBs^19,36-38^. The second DNA end-resection step consists of a 5’ to 3’ long-range resection mediated by two conserved^33^–^35^, separate pathways dependent on either the exonuclease Exo1 or the helicase-nuclease Sgs1 (the *S. pombe* Rqh1 and the mammalian BLM orthologue)-Dna2 ^27,28^. The long range-resection creates a longer 3’ tailed ssDNA coated by RPA, up to 2-4 kb^29^.

The nucleolytic processing of nascent strands at altered replication forks is central to resume DNA synthesis but uncontrolled resection is detrimental to genome stability^39^. There is a growing interest in understanding how ssDNA at forks occurs since ssDNA is a key activator of the DNA replication checkpoint, an anti-cancer therapeutic target^40^. Also, fork-degradation prevented by BRCA2 plays a pivotal role in the chemo-resistance of breast cancer cell lines^41^. Many of the factors required for DSBs resection are also involved in the end-resection of replication forks, such as MRN, DNA2 and CtIP^13,15,42-44^. However, how the steps of fork resection are orchestrated, compared to DSBs is poorly understood.

Here, we investigated the formation of ssDNA gaps using a model of terminally-arrested and unbroken forks. The resection of newly replicated strands occurs in two-steps. An initial resection mediated by MRN/Ctp1 generates short ssDNA gaps of ~ 110 bp in size which are obligatory to promote Rad51-mediated fork restart. A long-range resection mediated by Exo1, but not Rqh1, generates larger ssDNA gaps which are dispensable to replication restart. Unexpectedly, despite the absence of DSBs, we found that MRN/Ctp1 removes Ku from terminally-arrested forks, allowing long-range resection to occur. The lack of Ku results in a slower replication fork restart, characterized by an extensive resection but surprisingly a reduced amount of RPA loading. Our data are consistent with dysfunctional forks undergoing fork reversal, which provides a single DNA end for Ku binding. We uncover an unprecedented role for Ku in orchestrating the resection of terminally-arrested, in the absence of DSBs, and in fine-tuning replication restart.

## Results

### A genetic assay to investigate replication fork-resection and restart at a conditional RFB in fission yeast

We exploited a previously described conditional RFB, named *RTS1*, to block a replisome in a polar manner at a specific locus^45^ (**Fig. 1a**). The blockage of the replication fork is mediated by the *RTS1*-bound protein Rtf1 whose expression is controlled through the *nmt41* promoter repressed in the presence of thiamine. The *RTS1*-RFB has been integrated at the natural *ura4* locus, at which most forks travel from the centromere-proximal origin towards the telomere. 16 hours after thiamine removal, > 90% of forks travelling in the main replication direction are blocked at the *RTS1*-RFB resulting in dysfunctional forks. These terminally-arrested forks are either restarted by HR or rescued by the progression of opposite forks^17^. HR-mediated replication restart occurs in 20 minutes and is initiated by a ssDNA gap, not a DSB, on which RPA, Rad52 and Rad51 are loaded^46-48^.

**Figure 1.**
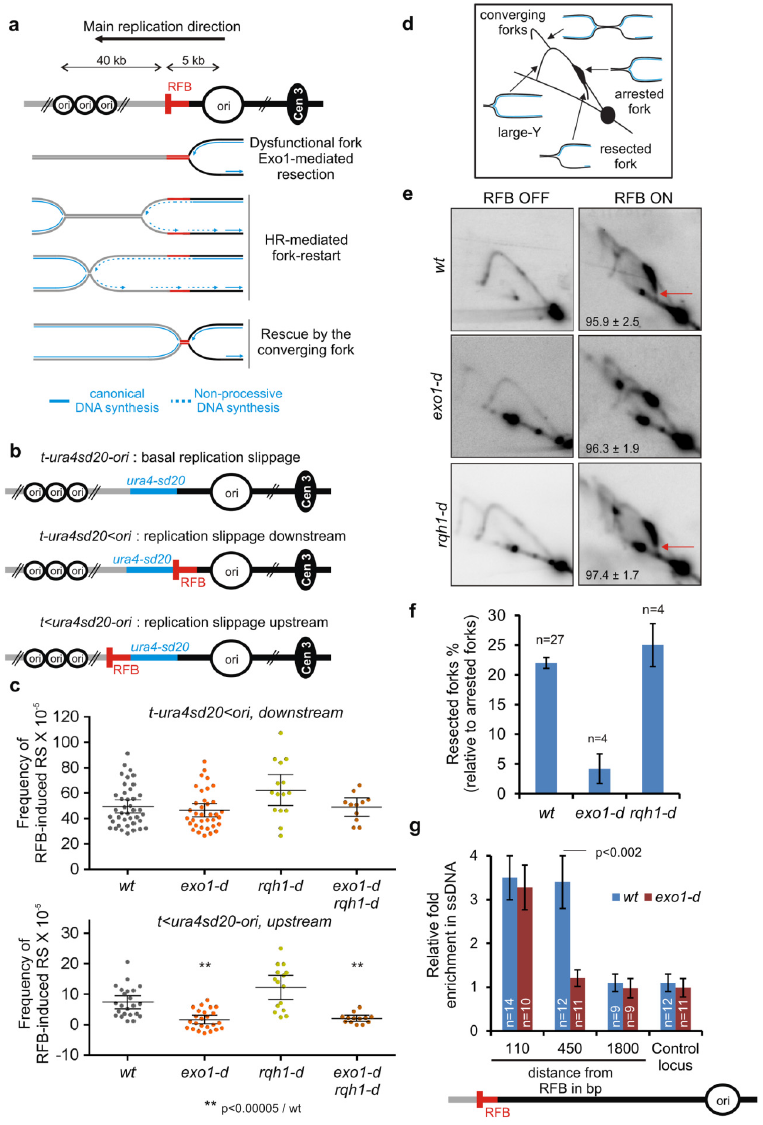
Long-range resection of terminally-arrested forks relies on Exo1 but not Rqh1. **(a)** Scheme of the *RTS1*-RFB (red bar) integrated 5 kb away from a strong replication origin (ori). “Cen3” indicates position of centromere. Distances between telomere-proximal replication origins and the *RTS1*-RFB are indicated. Upon Rtf1 expression, forks traveling from the centromere toward the telomere are blocked in a polar manner. **(b)** Diagrams of constructs containing the reporter gene *ura4-sd20*, associated or not to the *RTS1*-RFB (see text and Supplementary Fig. 1). **(c)** Frequency of upstream and downstream RFB-induced Replication Slippage (RS). Each dot represents one sample. Bars indicate mean values ± 95 % confidence interval (CI). Statistics were calculated using Mann and Whitney U test (See Supplementary excel file 1). **(d)** Scheme of replication intermediates (RI) analysed by neutral-neutral 2DGE of the *AseI* restriction fragment. **(e)** Representative RI analysis by 2DGE in RFB ON and RFB OFF conditions. Red arrows indicate terminally-arrested forks containing nascent strands undergoing resection. Numbers indicate % of blocked forks ± standard deviation (SD). **(f)** Quantification of resected forks (tail signal), relative to the intensity of terminally-arrested forks. Values are means of *n* samples from independent experiments ± 95% CI. **(g)** Relative enrichment of ssDNA formed upstream from the *RTS1*-RFB (see Methods). Data show the fold enrichment in ssDNA in the RFB ON relative to the RFB OFF condition. A locus located on chromosome II is used as control. Values are means of n samples from independent experiments ± standard error of the mean (SEM). Statistics were calculated using Mann and Whitney U test.

We previously reported that HR-mediated fork restart is associated with a non-processive DNA synthesis, liable to replication slippage (RS)^49^. Both strands of restarted forks are synthetized by Polymerase delta, reflecting a non-canonical replisome likely insensitive to the RFB^50^. We developed a reporter assay consisting of an inactivated allele of *ura4, ura4sd20*, which contains a 20 nt duplication flanked by 5 bp of micro-homology^49^. When the *ura4sd20* allele is replicated by a restarted fork, the non-processive DNA synthesis undergoes RS resulting in the deletion of the duplication and the restoration of a functional *ura4*^+^ gene. The *ura4sd20* allele was inserted either downstream or upstream from the *RTS1*-RFB to generate the construct *t-ura4sd20<ori* or *t<ura4sd20-ori* (t for telomere, < for the *RTS1*-RFB and its polarity), respectively (**F*ig*. 1b**). We showed that RS occurring upstream and downstream from the *RTS1*-RFB are both consequences of restarted forks^47^. To monitor the basal level of RS in different genetic backgrounds, we also generated a *t-ura4sd20-ori* construct, devoid of the *RTS1*-RFB.

The frequency of downstream RS was increased by 23 fold upon expression of Rtf1 (*t-ura4sd20*<ori, compare Rft1 repressed to expressed situation, **Supplementary Fig. 1a**) and by 17 fold compared to the strain devoid of RFB (*t-ura4sd20-ori*, **Supplementary Fig. 1a**). In *rad51* or *rad52 deleted strains*, the leakiness of Rft1 repression was more obvious as the RS frequency was slightly higher when Rft1 was repressed compared to the strain devoid of RFB (**Supplementary Fig. 1a**, compare Rtf1 expression: -, *RTS1*: -, RFB activity: - with Rtf1 expression: -, *RTS1*: +, RFB: L). Thus, to obtain the true occurrence of RS by the *RTS1*-RFB, independently of the genetic background, we subtracted the RS frequency of the strain devoid of RFB from the frequency of the strain containing *the t-ura4sd20<ori* construct, upon expression of Rtf1 (**Supplementary Fig. 1a**, red arrows).The frequency of RS induced by the *RTS1*-RFB was of 49.6, 9.8 and 10.8×10^−5^, in *wt, rad51-d* and *rad52-d* strain, respectively (**Supplementary Fig. 1b**). We estimated that the lack of Rad51 or Rad52 results in 80 % of forks irreversibly and terminally arrested at the RFB.

The frequency of upstream RS was increased by 3 fold in *wt* cells (compared to the strain devoid of RFB) and this induction was abolished in *rad52-d* and *rad51-d* cells (**Supplementary Fig. 1c-e**). Indeed, we reported that the DNA synthesis associated to Rad51-mediated fork restart occurs occasionally upstream from the initial site of fork arrest, as a consequence of ssDNA gap formation by Exo1^47^.

Thus, the RS assays associated to the *RTS1*-RFB allows the quantification of replication restart efficiency and an indirect monitoring of fork-resection.

### The long-range resection is performed by Exo1, but not Rqh1, and is dispensable to replication restart

HR-mediated fork restart at the *RTS1*-RFB is initiated by ssDNA gaps of ~ 1kb in size formed upstream from terminally-arrested forks^47^. Although Exo1 is the main nuclease responsible for the formation of these gaps, the lack of Exo1 does not impair the efficiency of replication restart, suggesting that additional nucleases are likely involved. Consistent with this, upstream RFB-induced RS was abolished in *exo1-d* cells, whereas downstream RFB-induced RS was not affected (**Fig. 1c**). The DSB long-range resection is mediated by two independent parallel pathways: Exo1 and Rqh1^Sgs^1^/BLM^/Dna2^27,28^. We applied the RS assays to *rqh1-d* cells. The lack of Rqh1 did not affect upstream and downstream RFB-induced RS, even when *exo1* was deleted, indicating that Rqh1 is not required for fork-resection and restart (**Fig. 1c**).

We recently reported a novel method to monitor fork-resection by bi-dimensional gel electrophoresis^51^ (2DGE). We identified an intermediate emanating from the fork arrest signal and descending towards the linear arc, indicative of a loss of mass and shape (**Fig. 1d-e**, see red arrow). Alkaline 2DGE showed that this tail signal corresponds to terminally-arrested forks containing newly replicated strands undergoing resection^51^. Consistent with this, the tail signal was abolished in *exo1-d* (**Fig. 1e-f**). We analysed fork-resection by 2DGE in *rqh1-d* cells. The tail signal was unaffected in *rqh1-d* cells compared to *wt* cells, confirming that, in contrast to DSBs, Rqh1 is not part of a long-range resection pathway of terminally-arrested forks (**Fig. 1e-f**).

To get a deeper analysis of ssDNA length generated upstream from the *RTS1*-RFB, we recently developed a qPCR assay to directly monitor ssDNA, based on ssDNA being refractory to restriction digestion^51,52^. As previously reported, in *wt* cells, ssDNA was enriched 110 bp and 450 bp upstream from the *RTS1*-RFB but was undetectable at 1.8 Kb from the RFB. Yet, in the absence of Exo1, ssDNA was still enriched at 110 bp but not at 450 bp (**Fig. 1g**). These data hint at the presence of additional nucleases acting on terminally-arrested forks to generate short ssDNA gap and subsequent replication restart.

### MRN/Ctp1 promotes fork-resection and restart, independently of the Mre11 nuclease activity

We investigated the role of Rad50 and Ctp1 in fork-resection and restart. Compared to *wt* cells, downstream RFB-induced RS was decreased by 1.8 and 3.4 fold in *ctp1-d* and *rad50-d* cells, respectively (**Fig. 2a**, upper panel). The *ctp1-d rad50-d* double mutant showed a defect similar to each single mutant, showing that MRN and Ctp1 act in the same pathway of HR-mediated fork restart. We estimated that the lack of MRN/Ctp1 results in ~ 70 % of forks irreversibly terminally-arrested at the *RTS1*-RFB.

**Figure 2.**
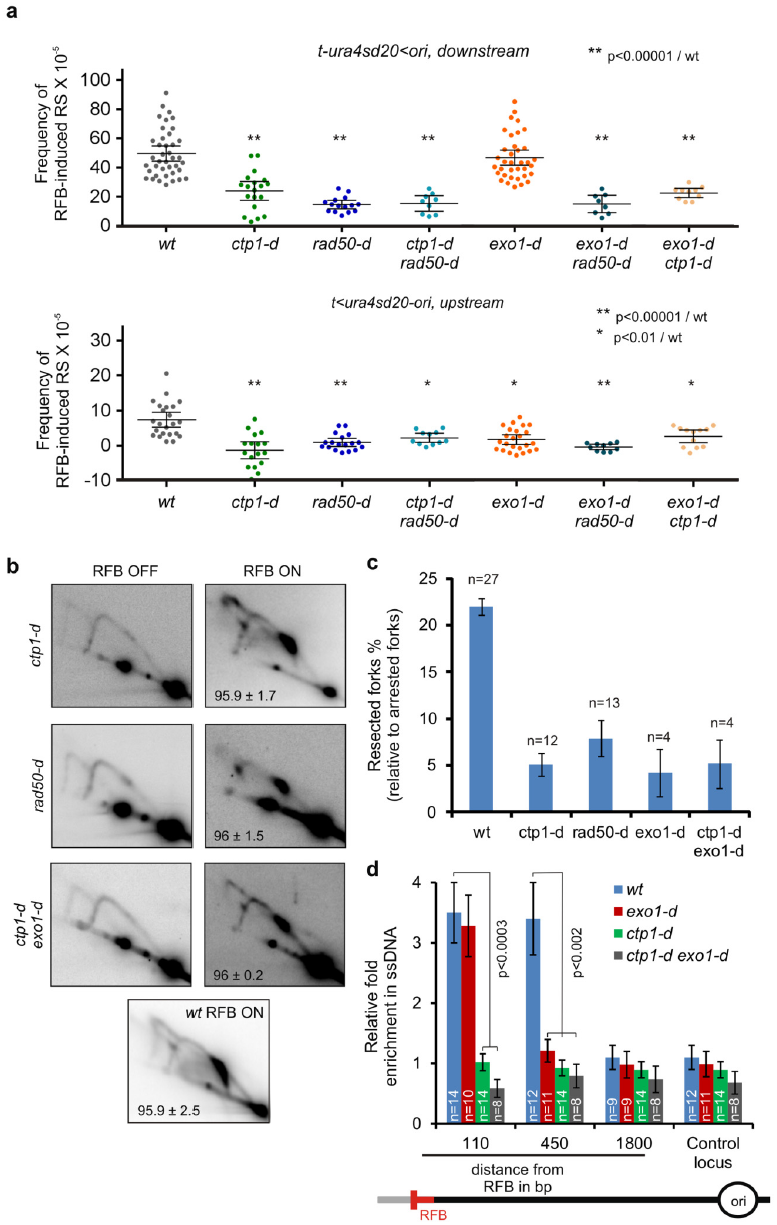
MRN/Ctp1-mediated initial fork resection is essential to replication restart and primes long-range resection by Exo1. **(a)** Frequency of RFB-induced RS downstream (top panel, *t-ura4sd20<ori*) and upstream (bottom panel, *t<ura4sd20-ori*) as described on Fig. 1c. **(b)** Representative RI analysis by 2DGE in indicated strains upon activation (RFB ON) or not (RFB OFF) of the *RTS1*-RFB, as described on Fig. 1e. n indicates the number of samples. **(c)** Quantification of forks undergoing resection (“tail signal”), relative to the intensity of terminally-arrested forks, as described on Fig. 1f. n indicates the number of samples. **(d)** Relative enrichment of ssDNA formed upstream from the *RTS1*-RFB, as described on Fig. 1g. n indicates the number of samples. Statistics were calculated using the non-parametric Mann and Whitney U test. See Supplementary Fig. 2 for the analysis of the mre11 nuclease dead mutant.

As observed in *exo1-d* cells, upstream RFB-induced RS was similarly abolished in *ctp1-d, rad50-d* and, *ctp1-d rad50-d* double mutant, suggesting that MRN/Ctp1 may act in fork-resection to ensure efficient DNA synthesis resumption (**Fig. 2a**, bottom panel). Consistent with this, analysis of fork-resection by 2DGE showed that the level of resected forks was severely and similarly decreased in either *rad50-d* or *ctp1-d cells* (**Fig. 2b-c**). We concluded that MRN/Ctp1 act together to promote fork-resection and subsequent fork-restart.

We reported that Rad52 recruitment to the *RTS1*-RFB relies on Mre11, independently of its nuclease activity. We analysed RFB-induced RS and fork-resection by 2DGE in cells expressing a nuclease-dead form of Mre11 (Mre11-D65N), defective for both the endo- and exo-nuclease activities^53,54^. This mutant showed neither a defect in restarting replication forks nor a defect in resecting dysfunctional forks blocked at the *RTS1*-RFB (**Supplementary Fig. 2**). Thus, MRN/Ctp1-dependent fork-resection and restart requires an intact MRN complex but not the Mre11 nuclease activity.

### MRN/Ctp1 promotes short ssDNA gaps to prime Exo1-mediated long-range fork resection

In contrast to Exo1-mediated fork-resection, the MRN/Ctp1-dependent resection step is critical to ensure an efficient restart of DNA synthesis. We tested if MRN/Ctp1 acts upstream Exo1 in promoting the formation of ssDNA gaps. In the simultaneous absence of Exo1 and either Ctp1 or Rad50, upstream and downstream RFB-induced RS was decreased to the same extent as the single deletion of either *ctp1* or *rad50* (**Fig. 2a**). Fork-resection analysis by 2DGE revealed a similar defect in the single *ctp1-d* mutant than in the double *ctp1-d exo1-d* mutant (**Fig. 2b-c**). These data argue that the role of MRN/Ctp1 is not redundant with Exo1 function, and that MRN/Ctp1 acts upstream Exo1. To clarify the role of these factors in ssDNA gaps formation, we monitored ssDNA by qPCR in *ctp1-d* and *ctp1-d exo1-d* cells. ssDNA enrichment at both 110 bp and 450 bp was dependent on Ctp1. Thus, short and large ssDNA gaps are dependent on the MRN/Ctp1 pathway (**Fig. 2d**). Furthermore, Exo1 was no longer required for ssDNA formation at 110 bp in the absence of Ctp1. As shown for DSBs resection^27,28^, we propose that fork-resection is a two-step process: MRN/Ctp1 promotes the formation of short ssDNA gaps of ~ 110 bp in size which primes the Exo1-mediated long-range resection; the initial resection being critical for replication fork restart, but not the extensive one.

### Ku accumulates at terminally-arrested forks in the absence of MRN/Ctp1

Despite the absence of DSBs at the *RTS1*-RFB, we tested the role of Ku in the resection of terminally-arrested forks. The deletion of *pku70* on its own did not impact upstream RFB-induced RS, but rescued the defect observed in *rad50-d* and *ctp1-d* cells (**Fig. 3a**). To confirm that this rescue occurs at the step of fork-resection, we analysed resected-forks by 2DGE. The level of resected forks was fully restored in *ctp1-d pku70-d* and *rad50-d pku70-d* cells (**Fig. 3b-c**). Thus, the lack of Ku bypasses the requirement of Rad50/Ctp1-mediated initial resection of terminally-arrested forks.

**Figure 3.**
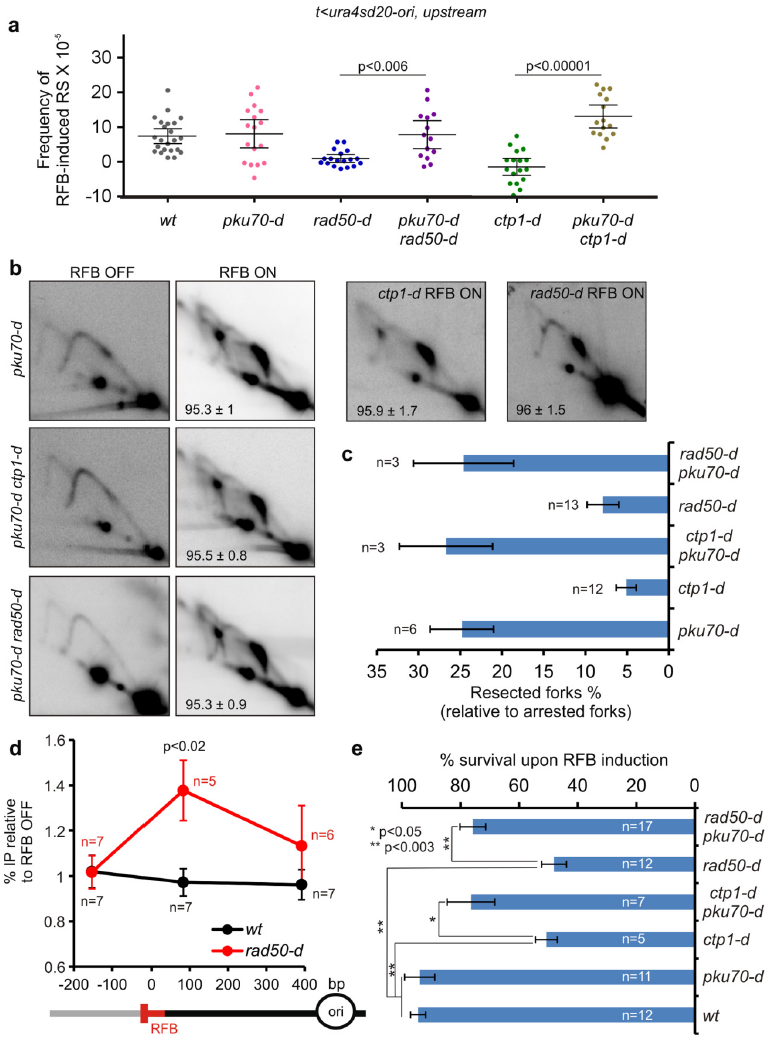
MRN/Ctp1 allows Ku eviction form terminally-arrested and unbroken forks. **(a)** Frequency of upstream RFB-induced RS, as described on Fig. 1c. **(b)** Representative RI analysis by 2DGE upon activation (RFB ON) or not (RFB OFF) of the *RTS1*-RFB, as described on Fig. 1e. Numbers indicate the % of forks blocked at the *RTS1*-RFB ± SD. N indicates the number of sample. **(c)** Quantification of forks undergoing resection (“tail signal”), relative to the intensity of terminally-arrested forks, as described on Fig. 1f. n indicates the number of samples. **(d)** Analysis of Ku recruitment to the *RTS1*-RFB by ChIP-qPCR in indicated strains. The fold-enrichment in the ON condition (RFB active) relative to the OFF condition (RFB inactive) is shown. Upstream and downstream distances from the RFB are indicated in base pairs (bp). The values are the means of at least five independent experiments ± SEM. n indicates the number of samples. Statistical analysis was performed using Mann-Whitney U tests. **(e)** Survival of the indicated strains upon activation of the barrier relative to OFF condition. Values are the means of at least 5 independent samples ± SEM. n indicates the number of samples. Statistics were calculated using the non-parametric Mann and Whitney U test. See Supplementary Fig. 3 for the analysis of cell sensitivity to CPT and MMS.

We investigated the recruitment of Pku70 to the *RTS1*-RFB by ChIP-qPCR. In *wt* cells, we were unable to detect any recruitment, which likely reflects that the binding of Ku to terminally-arrested forks is momentary, as observed to DSBs^36^ (**Fig. 3d**). In the absence of Rad50, Pku70 accumulated exclusively ~110bp upstream from the *RTS1*-RFB, suggesting that Ku binds near or at the expected 3-branched junction of the dysfunctional fork. Collectively, these data argue a role of MRN/Ctp1 in releasing Ku from terminally-arrested forks.

The lack of Ku suppresses the sensitivity of *ctp1-d* and *rad50-d* cells not only to DSBs-inducing agents but also to replication-blocking agents^19,22,23,36^. These data were interpreted as a role of Ku in binding replication-born DSBs formed in the vicinity of stressed forks. As reported, we found that the deletion of *pku70* rescued partially the sensitivity of *ctp1-d* and *rad50-d* cells to very low doses of camptothecin (CPT) and methylmethane sulfonate (MMS) (**Supplementary Fig. 3**). CPT is an inhibitor of topoisomerase I which slows down replication fork progression while MMS is an alkylated agent resulting in damaged replication forks. Both drugs do not induce detectable DSBs at very low doses^55,56^. We tested whether similar genetic interactions were observed upon induction of the *RTS1*-RFB. Both *ctp1-d* and *rad50-d* cells showed a loss of viability upon induction of the *RTS1*-RFB, that is rescued by deleting *pku70* (**Fig. 3e**). Altogether, these data support that the inability of MRN/Ctp1 to remove Ku from terminally-arrested forks is a lethal event.

### Ku orchestrates the initial and long-range resection of terminally-arrested forks

In the absence of Ku, MRN/Ctp1 is no longer needed to promote ssDNA gap formation. We tested whether fork-resection is dependent on Exo1 in the absence of Ku. Firstly, upstream RFB-induced RS observed in the *pku70-d ctp1-d* double mutant was dependent on Exo1 (**Fig. 4a**). Secondly, 2DGE analysis revealed that Exo1 is responsible for fork-resection occurring in *ctp1-d pku70-d* cells (**Fig. 4b-c**). Thirdly, as reported by Langerak et al.^36^, the suppressive effect of *pku70* deletion on *cpt1-d* cells sensitivity to CPT and MMS was dependent on Exo1 as the *pku70-d cpt1-d exo1-d* triple mutant exhibited the same sensitivity as the single *ctp1-d* mutant (**Supplementary Fig. 3**). These data establish that the bypass of initial fork-resection by the lack of Ku relies on Exo1. We propose that Ku has an inhibitory effect on the Exo1-mediated long-range fork-resection, requiring a MRN/Ctp1 relief, as proposed for DSBs and telomere resection.

**Figure 4.**
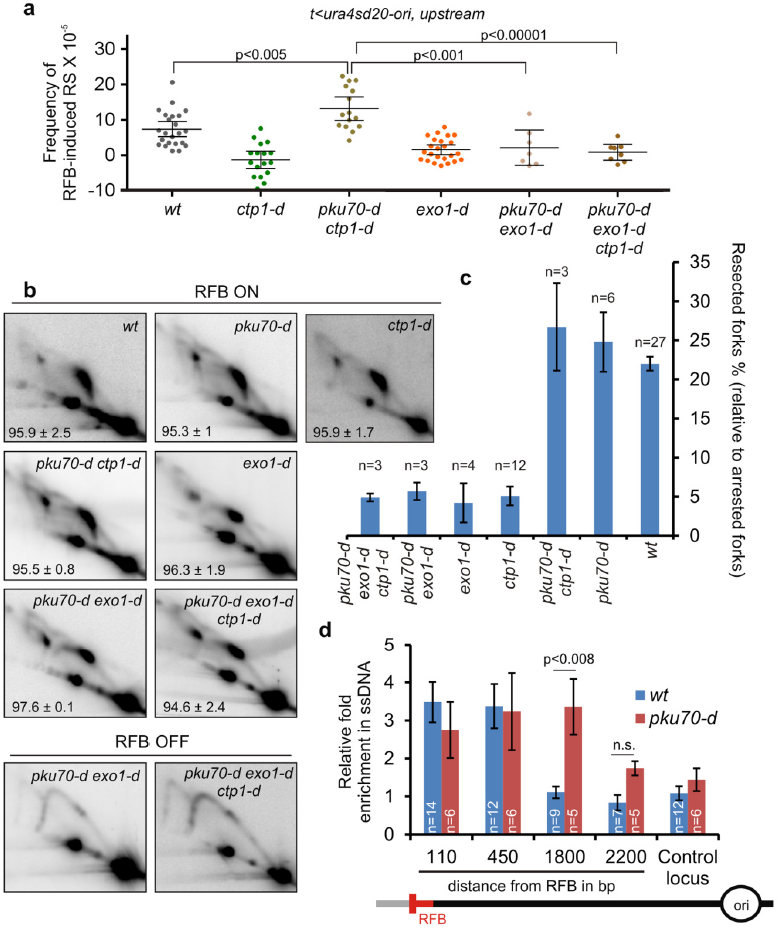
Ku orchestrates the two-step process of fork-resection. **(a)** Frequency of upstream RFB-induced RS, as described on Fig. 1c. **(b)** Representative RI analysis by 2DGE upon activation (RFB ON) or not (RFB OFF) of the *RTS1*-RFB, as described on Fig. 1e. **(c)** Quantification of forks undergoing resection (“tail signal”), relative to the intensity of terminally-arrested forks, as described on Fig. 1f. n indicates the number of samples. **(d)** Relative enrichment of ssDNA formed upstream from the *RTS1*-RFB in indicated strain. The data shown are the fold enrichment in ssDNA in the ON condition (RFB active) relative to the OFF condition (RFB inactive). A locus located on chromosome II is used as control. The values are the mean of at least three independent experiments ± standard error of the mean (SEM). n indicates the number of samples. Statistics were calculated using the non-parametric Mann and Whitney U test.

In the absence of Ku, terminally-arrested forks are no longer resected in a two-step manner and ssDNA gaps are directly formed by Exo1. Quantification of ssDNA by qPCR showed that ssDNA was significantly more abundant at 1.8 Kb upstream from the *RTS1*-RFB in *pku70-d* cells compared to *wt* cells (**Fig. 4d**). 2DGE analysis revealed that this extensive fork-resection relies on Exo1 (**Fig. 4b-c**). Thus, in the absence of Ku, terminally-arrested forks are excessively resected trough the Exo1-mediated long-range resection. We propose that Ku orchestrates the formation of ssDNA gaps by ensuring that fork-resection occurs in a two-step manner.

### The lack of Ku slows-down replication restart independently of NHEJ

To test whether the lack of Ku impairs HR-mediated fork restart, we applied the downstream RS assay to *pku70-d* cells. RFB-induced RS was decreased by 2.2 fold compared to *wt*, indicating that ~50 % of forks are difficult to restart (**Fig. 5a**). This cannot be explained by a lower expression of Rad51, Rad52 and RPA (**Supplementary Fig. 4a-b**). We tested a possible involvement of Ligase 4, and found no defect in upstream and downstream RFB-induced RS in *lig4-d* cells, showing that Ligase 4 is neither required for fork-resection nor restart (**Supplementary Fig. 4c**). These data suggest that Ku is involved in HR-mediated replication restart, independently of NHEJ.

**Figure 5.**
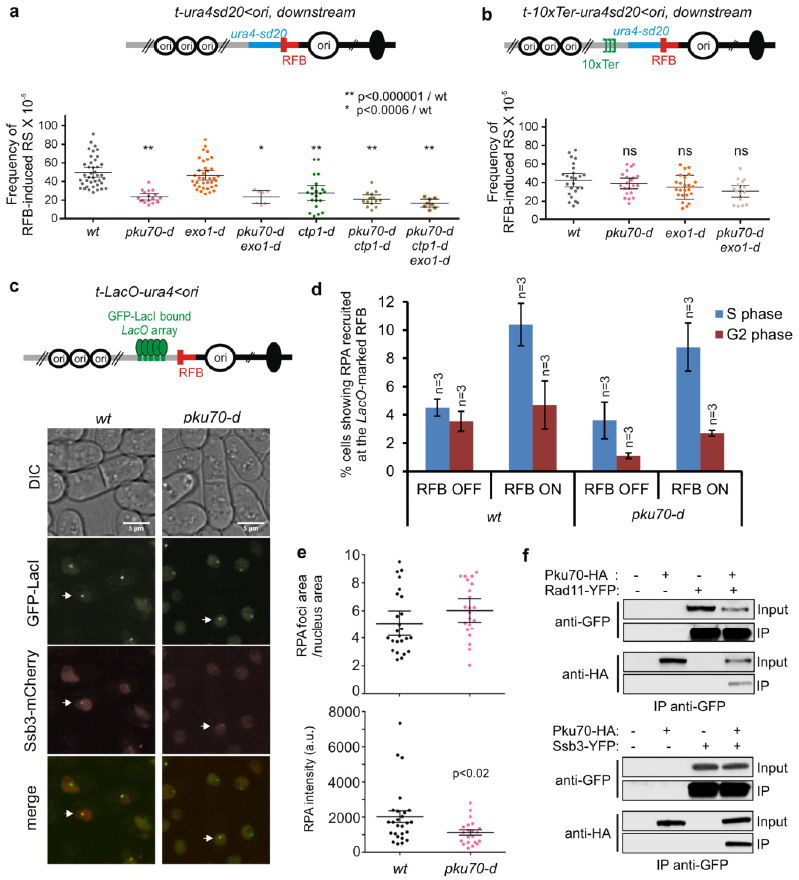
Ku fine-tunes HR-mediated fork restart and RPA loading onto ssDNA at terminally arrested forks. **(a)** Frequency of downstream RFB-induced RS, as described on Fig. 1c. **(b)** Frequency of downstream RFB-induced RS using the *t-10xTer-ura4sd20<ori* construct at which the progression of the converging fork is delayed. Each dot represents one sample. Bars indicate mean values ± 95 % CI. Statistics were calculated using the Mann and Whitney U test. See Supplementary Fig. 4 for Hr factors expression and analysis of the lig4-d mutant. **(c)** Top panel: scheme of the *LacO*-marked *RTS1*-RFB. Bottom panel: representative images showing RPA (labelled with Ssb3-mCherry) recruitment to the *LacO*-marked RFB. Of note, to avoid biases toward a random localization of GFP-LacI and RPA foci, cells with ≥ 3 RPA foci were excluded from the analysis. **(d)** % of cells showing RPA recruitment to the *LacO*-marked RFB, according to cell cycle phase. G2 and S-phase cells are mononucleated cells and bi-nucleated cells with a septum, respectively. Values are means of at least *n* independent experiments ± SD. **(e)** Area and intensity of RPA foci recruited to the *LacO*-marked RFB. Each dot represents one RPA foci in S phase upon RFB activation. Bars indicate the mean value ± 95% CI. Statistics were calculated using the non-parametric Mann and Whitney U test. **(f)** Co-immunoprecipitation of Pku70-HA with two subunits of RPA, Rad11 and Ssb3, fused to the YFP epitope. Experiments were performed in the presence of benzonase. For a longer exposure, see Supplementary Fig. 5.

We asked whether excessive fork-resection impairs replication restart. We analysed downstream RFB-induced RS in the *pku70-d exo1-d* double mutant in which long-range resection is abolished, and the *pku70-d ctp1-d*, and the *pku70-d rad50-d* double mutants, in which long-range resection still occurs (**Fig. 5 a-b, Supplementary Fig. 4d**). The three strains behave as the single *pku70-d* strain, with a ~ 2 fold reduction in RFB-induced RS compared to *wt* (**Fig. 5a**). The *pku70-d ctp1-d exo1-d* triple mutant showed up to 3 fold reduction in RFB-induced RS (**Fig. 5a**). Thus, the lack of Ku cripples HR-mediated fork restart whatever the extent of fork resection and the length of ssDNA gaps.

We further investigated the requirement of Ku in fork-restart at the *RTS1*-RFB. We employed a strain in which fork convergence from the distal side is minimized by the presence of 10 repeats of the TER2/TER3 rDNA rRFBs (**Fig. 5b**). Unlike the *RTS1*-RFB, TER2/TER3 slows down fork progression without inducing terminally-arrested forks and recruitment of HR factors^46^. Delaying the arrival of opposite forks allows more time for the process of HR-mediated fork-restart to occur at the *RTS1*-RFB^50^. We monitored the level of downstream RFB-induced RS in *wt* and *pku70-d* and observed no significant differences, even when the Exo1-long range resection was abolished (**Fig. 5b**). Thus, the defect in RFB-induced RS is rescued by delaying the arrival of converging forks. This suggests that the defect is a consequence of a slow HR-mediated fork restart process rather than an inability to restart replication-forks in the absence of Ku.

### Ku fine-tunes RPA-coated ssDNA and replication restart

We investigated the dynamics of RPA recruitment to the *RTS1*-RFB by fluorescence-based imaging in living cells. We employed a strain in which a GFP-LacI-bound *LacO* array was integrated closed to the the *RTS1*-RFB, and expressing the RPA subunit Ssb3 fused to mCherry^51^ (**Fig. 5c**). In both *wt* and *pku70-d* strains, we observed a similar increase in S-phase cells showing RPA recruitment to the *LacO*-marked RFB (**Fig 5c-d**). Such recruitment did not occur in G2 cells. Thus, RPA is recruited to resected forks in S-phase cells and then evicted once arrested forks have been restarted or rescued by converging forks. Thus, despite slowing down of fork restart in the absence of Ku, this does not result in an accumulation of resected and arrested forks in G2 cells, suggesting that dysfunctional forks are ultimately rescued by the progression of opposite forks.

When performing cell imaging, we noticed that RPA foci were less bright in the absence of Ku. We quantified the area and intensity of RPA foci recruited to the *LacO*-marked RFB. We found that the area was not affected, whereas the intensity of RPA foci was decreased by half in *pku70-d* cells compared to *wt* (**Fig. 5e**). These observations cannot be explained by a reduced amount of ssDNA in the absence of Ku, as ssDNA gaps are ~ twice longer than in *wt* cells (**Fig. 4d**). These data suggest a role for Ku in recruiting RPA onto ssDNA at terminally-arrested forks. In support of this, we observed that Pku70-HA co-immunoprecipitates with two RPA subunits, Rad11-YFP and Ssb3-YFP, from protein extracts treated with benzonase. Thus, as observed in budding yeast, Ku interacts with RPA, independently of DNA and RNA^57^ (**Fig. 5f and supplementary Fig. 5**). Altogether, these data indicate a role of Ku in ensuring RPA-coated ssDNA gaps at terminally-arrested forks to fine-tune HR-mediated replication restart.

## Discussion

Rad51-dependent processing of replication forks can occur independently of DSBs^14,17^. We investigated the resection step of terminally-arrested, unbroken, forks that allows the exposure of ssDNA, subsequent recruitment of RPA, and Rad51-mediated fork-restart^47^. We made unexpected findings. First, the two-step model of DSB resection applies to unbroken forks. Second, the initial fork-resection includes Ku eviction, suggesting that dysfunctional forks undergo fork-reversal, providing a single dsDNA end for Ku binding. Third, the lack of Ku impairs efficient ssDNA coating by RPA and slows down HR-mediated fork restart, independently of NHEJ.

### Ku is recruited to terminally-arrested and unbroken forks

Our genetic data establish that Ku inhibits Exo1-mediated long range resection of terminally-arrested forks. MRN/Ctp1 counteracts this inhibition. Ku binds dsDNA ends with high affinity, and with poor affinity for ssDNA^58^. Despite the lack of DSBs, we demonstrate that Ku is recruited to dysfunctional and unbroken forks. These surprising findings favor a model in which terminally-arrested forks undergo fork-reversal^55^ (**Fig. 6**). Reversed forks are DNA structures in which newly replicated strands are annealed together, exposing a single dsDNA end for Ku binding. Fork reversal was shown, in mammals, to occur in response to various replication stresses, such as low doses of CPT and MMS, even in absence of replication-born DSBs^14^. Ku enrichment was only detected in the vicinity of the expected 3-branched junction, and not further upstream of the *RTS1*-RFB, suggesting little or no sliding activity of Ku on the reversed arm.

**Figure 6.**
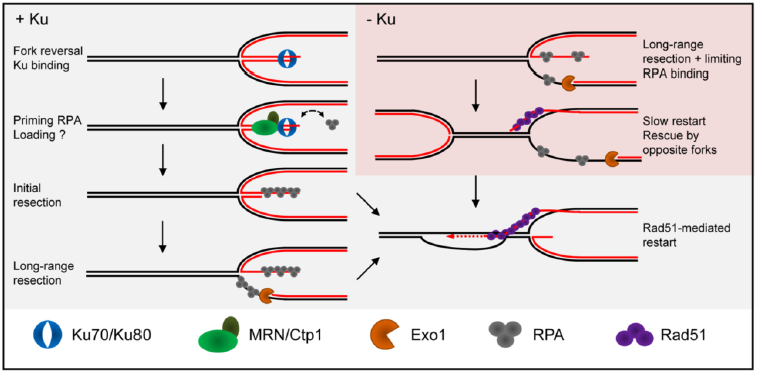
Model of Ku orchestrating the resection of terminally arrested fork to fine-tune Rad51-mediated replication restart. In *wt* cells, dysfunctional forks undergo fork reversal providing a single DNA end for Ku binding. Through physical interactions between Ku and RPA, Ku recruitment to replication fork primes for RPA loading onto ssDNA. MRN/Ctp1 allows Ku removal, generating a 110 bp sized ssDNA on which Rad51 nucleates to promote strand invasion within the restored parental duplex. In the absence of Ku, DNA end of reversed fork is directly resected by Exo1, resulting in extensive fork-resection and impaired RPA recruitment, which in turns slows down Rad51-mediated fork restart.

Ku is involved in the repair of replication-born DSB, either generated by defect in processing Okazaki fragments, CPT-induced, or genetically engineered^19,22-24,36^. The mammalian Ku associates with nascent strands after CPT treatment^59^ and fission yeast Ku is recruited to the rDNA RFB to stabilize blocked forks^26^. While these data are consistent with Ku being recruited to one ended DSB formed in the vicinity of replication forks, our finding suggest an alternative route to Ku recruitment, independently of DSBs. We propose that the DNA end of reversed fork is recognized and processed as a DSB, despite the absence of detectable DNA breakage (**Fig. 6**).

### MRN/Ctp1 ensures Ku removal to initiate fork-resection and restart

MRN and Ctp1 act together to expose ssDNA at terminally-arrested forks. The lack of MRN/Ctp1 results in Ku persisting at replication forks. Possibly, MRN/Ctp1 initiates fork-resection to create a substrate less favorable to Ku binding^19^. However, the lack of Ku impacts fork-resection and restart, suggesting that Ku binds terminally-arrested forks even if the MRN/Ctp1 pathway is functional. Thus, we propose that Ku is recruited early, likely on the reversed arm, and is then quickly evicted by MRN/Ctp1.

MRN has a catalytic and structural role in DSBs resection^60^. As observed for the resection of “clean” DSBs^32,36^, the Mre11 nuclease activity is dispensable to fork-resection and restart. The tetrameric form of Ctp1 has a scaffolding role in DSBs resection, through DNA binding and bridging activities^61^. Purified Ctp1 shows a binding preference for branched structure containing dsDNA and ssDNA, but no apparent nucleolytic activity was detected *in vitro*, in contrast to Sae2 and CtIP^61^. We propose that MRN/Ctp1 functions in initiating fork-resection and Ku eviction involves a structural rather than a nucleolytic role. Possibly, MRN/Ctp1 may recruit additional nucleases which remain to identify.

MRN/Ctp1 and their homologues counteract the accumulation of Ku at one ended DSB to promote Exo1-dependent resection^23,36,38^. The persistence of Ku on DNA end impairs the recruitment of yeast RPA and mammalian Rad51 without affecting end-resection^24,36^. In fission yeast, a single broken fork is lethal in *ctp1-d* cells^36^. Here, we establish a single terminally-arrested fork is lethal in the absence of either MRN or Ctp1. This lethality is in part caused by the binding of Ku to dysfunctional forks. Thus, we propose that Ku eviction from terminally-arrested forks by MRN/Ctp1 is an essential step for subsequent replication restart and cell viability.

### Ku coordinates the two-step process of fork-resection and fine-tunes Rad51-mediated fork restart

Together with our previous work, our data establish that the resection of terminally-arrested and unbroken forks occurs in two-steps which are coordinated by Ku. The initial MRN/Ctp1-dependent resection promotes Ku eviction and generates ~110 bp of ssDNA, essential to HR factors’ recruitment and subsequent fork-restart^47^. The second step is a long-range resection, mediated by Exo1, but not Rqh1, generating up to 1kb of ssDNA which is not strictly required to the resumption of DNA synthesis. Thus, a limited amount of ssDNA is sufficient to promote Rad51-mediated fork restart whereas long-range resection may reinforce checkpoint activation.

In the absence of Ku, MRN/Ctp1 is no longer required to initiate fork-resection, which then relies only on Exo1. As a consequence, terminally-arrested forks are extensively resected with an accumulation of ~2Kb of ssDNA. This supports that Exo1 inhibition by Ku takes place at the initial resection step even with a functional MRN/Ctp1 pathway. Remarkably, the initial resection of DSBs and telomeres is increased in the absence of Ku^21,38^. Thus, Ku plays an important role in ensuring that fork-resection occurs in two-steps to avoid unnecessary extensive resection, which can be detrimental to genome stability.

Previous works have proposed a NHEJ-independent role of Ku at replication-born DSBs, to channel repair towards the HR pathway ^22,23,36^. Fission yeast genetics indicate a role for Ku in the recovery from replication stress and stabilizing replication forks^25,26^. An important finding we made is that the lack of Ku slows down Rad51-mediated fork restart, accompanied with an extensive fork resection and a reduced amount of RPA-coated ssDNA. These data contrast with Ku acting as a barrier against RPA loading onto ssDNA during DSBs repair^36^, suggesting replicative-specific functions for Ku. RPA is a critical factor to control end-resection, stability of ssDNA and subsequent recruitment of HR factors^62^. As observed in budding yeast^57^, we report physical interactions between Ku and RPA, suggesting that Ku facilitates RPA loading onto ssDNA. We propose that Ku fine-tunes Rad51-mediated fork restart by priming RPA loading onto ssDNA and subsequent HR factors’ recruitment.

Ku allows the recruitment of downstream NHEJ factors such as Ligase 4 to promote DSBs ligation. We found no role for Ligase 4 in promoting fork-resection and restart. Our data rather suggest that NHEJ inhibition enhances HR-mediated replication restart. Given the potential deleterious outcomes of error-prone NHEJ events on genome stability, it is possible that the recruitment of additional NHEJ factors is prevented to avoid unwanted ligation of the reversed arm at terminally-arrested forks.

Recent reports indicate that ssDNA stabilization by RPA influences repair pathway choice at DSB between BIR and microhomology-mediated end joining (MMEJ)^63,64^. MMEJ is an alternative NHEJ mechanism, independent of Ku and ligase IV, that promotes DSB repair toward chromosomal rearrangement. We propose that the lack of Ku, resulting in reduced RPA-coated ssDNA at dysfunctional forks, may favor MMEJ events.

## Online Method

### Standards yeast genetics

Yeast strains used in this work are listed in Supplementary Table 1. Gene deletion and gene tagging were performed by classical and molecular genetics techniques ^65^. Strains containing the replication *RTS1*-RFB were grown in supplemented EMM-glutamate media containing thiamine at 60°M. To induce the *RTS1*-RFB, cells were washed twice in water and grown in supplemented EMM-glutamate media containing thiamine (Rtf1 repressed, RFB OFF condition) or not (Rtf1 expressed, RFB ON condition) for 24 hours.

### Analysis of replication intermediates by 2DGE

Replication intermediates were analyzed by 2DGE as described in Ait Saada et al.^51^. Exponentially growing cells (2.5×10^9^) were harvested with 0.1% sodium azide and frozen EDTA (80mM final concentration). Genomic DNA was crosslinked by adding trimethyl psoralen (0.01mg/ml, TMP, Sigma, 3902-71-4) to the cell suspensions, for 5min in the dark. Then, cells were exposed to UV-A (365nm) for 90 seconds at a flow of 50mW/cm^2^. Cells were lysed with 0.625mg/ml lysing enzyme (Sigma, L1412) and 0.5mg/ml zymolyase 100T (Amsbio, 120493-1). The resulting spheroplasts were then embedded into 1% low melting agarose (InCert Agarose, Lonza) plugs, and then incubated overnight at 55°C in a digestion buffer containing 1mg/ml of proteinase K (Euromedex EU0090) and then washed and stored in TE (50mM Tris, 10mM EDTA) at 4ଌ. DNA digestion was performed with 60 units per plug of the restriction enzyme *Ase*I and equilibrated at 0.3M NaCl. Replication intermediates were enriched using BND cellulose columns (Sigma, B6385) as described in Lambert *et al.*, 2010. RIs were migrated in 0.35% agarose gel in TBE for the first dimension. The second dimension was migrated in 0.9% agarose gel-TBE supplemented with EtBr^66^. DNA was transferred to a nylon membrane in 10X SSC. Membranes were incubated with a ^32^P radio-labeled *ura4* probe, and RIs were detected using phosphor-imager software (Typhoon-trio) and quantified with ImageQuantTL.

### Live cell imaging

Cell preparation was done as described in Ait Saada et al.^51^. Cells were grown in supplemented filtered EMM-glutamate, washed twice and resuspended in fresh filtered media without supplements. A drop of 1-2μl of exponentially growing cultures was placed on a well of a microscope slide (Thermo Scientific) coated with 1.4% ultrapure agarose (Invitogen, Ref: 16500-500) prepared in filtered EMM. Acquisition was performed using an automated spinning disc confocal microscope (Nikon Ti Eclipse inverted) equipped with an ORCA Flash 4.0 camera, a temperature control box set at 30°C and a 100x oil immersion objective with a numerical aperture of 1.49. Z-stack image acquisition was performed using METAMORPH software, and the analysis of the resulting images was performed using ImageJ software. Image acquisition and analysis were performed on workstations of the PICT-IBiSA Orsay Imaging facility of Institut Curie. Ssb3-mCherry and GFP-LacI foci merging/touching was analyzed taking into account cell cycle phase and RFB activation (OFF and ON). “Foci merging/touching” refers to foci that partially or completely overlap, or to joint foci. The area and of Ssb3-mCherry foci that merge with GFP-LacI foci in S phase cells in ON condition was normalized by the total area of the nucleus. Ssb3-mCherry foci intensity was measured using ImageJ (IntDen); to that value, the nucleus background fluorescence intensity was subtracted.

### DNA extraction and quantification of ssDNA by qPCR

DNA extraction was performed as described in Ait Saada et al.^51^. 1 to 2×10^8^ cells were mechanically broken by vortexing with glass beads (425-600μm, Sigma^®^). Genomic DNA was extracted by classical phenol/chloroform extraction. 5μg of DNA were digested (or not) with 100 units of the restriction enzyme *Mse*I. 30ng of digested and mock-digested DNA were amplified by qPCR using primers surrounding *Mse*I restriction site (primers are listed in supplementary table 2). ssDNA quantification was based on the work of Zierhut & Diffley^52^, using the formula:

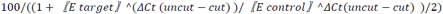

in which ΔCt is the difference between the threshold cycles of digested (cut) and undigested (uncut) DNA. *E* is amplification efficiency. A control locus (II-150) that does not contain *Mse*I restriction sites was used as a DNA loading control.

### Chromatin immunoprecipitation of Pku70-HA

Pku70-HA enrichment at *RTS1*-RFB was performed using strains expressing pku70-3xHA. ChIP experiments were performed as previously^36^, with the following modifications. Samples were crosslinked with 1% formaldehyde (Sigma, F-8775) for 15 minutes. Sonication was performed using a Diagenod Bioruptor at high setting for 8 cycles: 30 seconds ON + 30 seconds OFF. Immunopercipitation was performed using anti-HA antibody coupled to magnetic beads (Thermo Scientific, Pierce Anti-HA Magnetic Beads). Crosslink was reversed over night at 72°C. Samples were then incubated with Proteinase K and the DNA was purified using Qiagen PCR purification kit and eluted in 50 μl of elution buffer. The relative amount of DNA was quantified by qPCR (primers are listed in supplementary Table 2). Pku70-HA enrichment was normalized to an internal control locus (*ade6*). Data represent Ku enrichment in the ON condition relative to the OFF condition.

### Replication slippage assay with ura4-SD20 allele

Replication slippage using the *ura4-SD20* allele was performed as previously described^49^. 5-FOA resistant colonies were grown on uracil-containing plates with or without thiamine for 2 days at 30°C, and then inoculated in uracil-containing EMM for 24h. Cells were diluted and plated on YE plates (for survival counting) and on uracil-free plates containing thiamine to determine the reversion frequency. Colonies were counted after 5 to 7 days of incubation at 30°C. Statistics were performed using the non-parametric Mann and Whitney test. To avoid taking into account events that do not represent the behavior of each mutant (such as suppressor or other additional spontaneous mutations), outliers were not included in the statistical analysis or graphical representation. Outliers were calculated according to the formulas: Superior Outlier>1.5× (Q3-Q1)+Q3, and Inferior Outlier<1.5× (Q3-Q1)-Q1, where Q1 is the first quartile, and Q3 is the third quartile.

### Cell viability

Cell viability assays were performed as previously described^17^. Cells were grown on supplemented EMM without thiamine for 14 hours, then they were appropriately diluted and plated on EMM plates with (RFB OFF) or without thiamine (RFB ON). Colonies were counted after 5-7 days incubation at 30°C and viability was calculated as the ratio of colonies growing in ON condition relative to those growing in OFF condition.

### Co-immunopercipitation

5.10^8^ cells were harvested, washed in cold water and resuspended in 400 ml of EB buffer (50 mM HEPES High salt, 50mM KOAc pH 7.5, 5 mM EGTA, 1% triton X-100, 1 mM PMSF, and protease inhibitors). Cell lysis was performed with a Precellys homogenizer. The lysate was treated with 250mU/μl of benzonase for 30min. After centrifugation, the supernatant was recovered and an aliquot of 50 μl was saved as the INPUT control. To 300μl of lysate, 2 μl of anti-GFP (A11122 from Life Technologies) antibody were added and incubated for 1h30 at 4°C on a wheel. Then, 20μl of Dynabeads protein-G (Life Technologies) prewashed in PBS were added and then incubated at 4°C overnight. Beads were then washed twice 10 min in EB buffer before migration on acrylamide gel for analysis by Western blot. Ku70-HA, and Ssb3-YFP and Rad11-YFP were detected using anti-HA high affinity antibody (clone 3F10, Roche), and using anti-GFP antibody (Roche, clones 7.1 and 13.1), respectively. The supplementary Fig. 5 shows that Pku70-HA was slightly interacting in an unspecific way with anti-GFP antibody. However, the intensity of Pku70-HA in the IP fraction was highly increased in cells expressing SSb3-YFP or Rad11-YFP, showing that most interactions with Pku70-HA are specific.

## Acknowledgements

We thank Fuyuki Ishikawa for the gift of the Pku70-HA strain and Tony Carr for the *lig4-d* strain. We thank Vincent Geli for critical comments of the manuscript. We also thank the PICT-IBiSA@Orsay Imaging Facility of the Institut Curie. We thank Soraya Benazzouk and Yasmina Chekkal for their technical assistance. This study was supported by grants from the Institut Curie, the CNRS, the *Fondation ARC*, the *Fondation Ligue* (*comité Essone*), *l’Agence Nationale de la Recherche* ANR-14-CE10-0010-01, the *Institut National du Cancer* INCA 2013-1-PLBIO-14, and the *Fondation pour la Recherche Médicale* “Equipe FRM DEQ20160334889”. AAS, ATS, AC and II were funded by a French governmental fellowship, the Institut Curie international PhD program, *Association pour la Recherche sur le Cancer* (ARC), and the *Fondation pour la Recherche Medicale*, respectively. ATS was supported by a 4^th^ year PhD fellowship from *Fondation pour la Recherche Medicale* (FDT20160435131). The funders had no role in study design, data collection and analysis, the decision to publish, or preparation of the manuscript.

## Author contributions

A.T.S, A.A.S, I.I., M.N., K.F., J.H., and S.A.E.L. performed the experiments.

A.T.S A.A.S., I.I., and S.A.E.L. designed experiments and analyzed the data.

A.T.S and S.A.E.L wrote the paper.

## Supplementary Information

**Supplementary Figure 1:** A genetic assay to monitor HR-mediated replication restart and investigate fork-resection and the *RTS1*-RFB. (Related to Fig. 1)

**Supplementary Figure 2:** The nuclease activity of Mre11 is dispensable for fork-resection and restart. (Related to Fig. 2)

**Supplementary Figure 3:** The deletion of *pku70* partially rescues the sensitivity of ctp1-d and rad50-d cells to CPT and MMS, in an Exo1-dependent manner. (Related to Fig. 4)

**Supplementary Figure 4:** Ku promotes efficient replication restart, independently of NHEJ. (Related to Fig. 5)

**Supplementary Figure 5:** Higher exposure of the blot showed on the figure 5. (Related to Fig. 5)

**Supplementary table 1:** Strains used in this study

**Supplementary table 2:** Primers used in this study

**Supplementary excel file 1:** Statistics of RS frequency

